# Alternating lysis and lysogeny is a winning strategy in bacteriophages due to Parrondo’s Paradox

**DOI:** 10.1101/2021.03.07.434273

**Authors:** Kang Hao Cheong, Tao Wen, Sean Benler, Eugene V. Koonin

## Abstract

Temperate bacteriophages lyse or lysogenize the host cells depending on various parameters of infection, a key one being the host population density. However, the effect of different propensities of phages for lysis and lysogeny on phage fitness is an open problem. We explore a nonlinear dynamic evolution model of competition between two phages, one of which is disadvantaged in both the lytic and lysogenic phases. We show that the disadvantaged phage can win the competition by alternating between the lytic and lysogenic phases, each of which individually is a “loser”. This counter-intuitive result recapitulates Parrondo’s paradox in game theory, whereby individually losing strategies can combine to produce a winning outcome. The results suggest that evolution of phages optimizes the ratio between the lysis and lysogeny propensities rather than the phage reproduction rate in any individual phase. These findings are expected to broadly apply to the evolution of host-parasite interactions.

## Introduction

Bacteriophages outnumber all other reproducing biological entities in the biosphere combined, with an estimated instantaneous total of about 10^31^ virions across all biomes (Cobián Güemes et al., 2016). Bacteriophages reached this hyper-astronomical abundance using two basic strategies of infection that are traditionally classified as lytic and temperate. Lytic phages enter the host cells and immediately take over the cellular machinery to produce progeny virions, followed by a programmed burst of the cell (lysis) (Young, 2014) which releases progeny virions into the environment for subsequent rounds of infection. In contrast, temperate phages ‘decide’ to follow the lytic or lysogenic strategy at the onset of infection. In the lysogenic strategy, the phage DNA stably integrates into the host genome becoming a prophage that is inherited by the progeny during cell division and thus propagates with the host without lysis of the host cells. However, upon sensing an appropriate signal, such as DNA damage, a prophage ‘decides’ to end lysogeny and reproduce through the lytic pathway (Oppenheim et al., 2005; Ptashne, 2011). Given that an estimated 10^23^ infections of bacteria by bacteriophages occur every second (Hendrix, 2003), with profound effects on the global ecology as well as human health (Cobián Güemes et al., 2016; Liu et al., 2016; Manrique et al., 2017), the evolutionary processes that shape phage replication strategies of fundamental biological interest and importance.

The ability of temperate phages to decide between lysis or lysogeny has drawn considerable attention of theorists aimed at quantifying the conditions where one strategy prevails over the other or, in other words, deciphering the rules of phage lysis vs lysogeny decisions. Temperate viruses that choose lysogeny are effectively constrained by cellular binary fission, whereas lytic replication can produce bursts of hundreds and thousands of progeny virions from a single cell. A foundational theoretical study asked the question simply: why be temperate (Stewart and Levin, 1984)? The potential benefits of a non-lytic strategy are realized when host cell densities are too low to support lytic growth that would otherwise collapse one or both populations, and furthermore, the frequency of encounters of the phage particles released upon host lysis with uninfected host cells is low. In essence, lysogeny is advantageous at hard times (Stewart and Levin, 1984). Several recent formal model analyses agree that lysogeny is favored at low host cell density (Maslov and Sneppen, 2015; Wahl et al., 2019; Weitz et al., 2019). However, lysogeny also appears to be the dominant behavior at high host cell abundance (Knowles et al., 2016). The mechanisms driving viruses towards lysogeny at both low and high host cell abundances are not well understood, but differential cellular growth rates and viral adsorption rates appear to contribute (Luque and Silveira, 2020). Collectively, these studies underscore the importance of density-dependent dynamics for infection outcomes.

The paradigm for the decisions temperate phages make on the lytic or lysogenic pathway upon infection is *Escherichia* phage Lambda. Seminal work with Lambda has demonstrated that lysogeny is favored when multiple Lambda virions coinfect the same cell (Kourilsky, 1973). The standard interpretation is that coinfection is a proxy measurement for host cell density, driving Lambda towards lysogeny at low density (Herskowitz and Hagen, 1980). The genetic circuitry underlying Lambda’s “lysogenic response” has been meticulously dissected over decades of research (Casjens and Hendrix, 2015; Oppenheim et al., 2005; Ptashne, 2004) and continues to divulge new mechanistic determinants (Lee et al., 2018; St-Pierre and Endy, 2008; Trinh et al., 2017; Zeng et al., 2010). Directed evolution of Lambda yields mutants with different thresholds for switching from lysogeny to lysis (induction) and such heterogeneity has been observed in numerous Lambda-like phages (Deng et al., 2014; Murphy et al., 2008; Refardt and Rainey, 2010). Moreover, the vast genomic diversity of phages implies a commensurately diverse repertoire of lysis-lysogeny circuits, and experiments with phages unrelated to Lambda have revealed a variety of ways evolution constructed these genetic switches (Broussard et al., 2013; Erez et al., 2017; Ravin et al., 1999). How the different propensities for lysis or lysogeny impact phage fitness at different host cell densities, and in particular, when in competition with other phages, remains an open problem.

Inspired by the previous theoretical and experimental studies, we propose a population model to investigate the competition between two phages that differ in their rates of establishing lysogeny based on host cell density. In this model, the first phage (*P*_1_) has a higher mortality rate and lower reproduction rate during both lysis and lysogeny compared to the second, competing phage *P*_2_. From a game-theoretical perspective, *P*_1_ is burdened by two losing strategies. Unexpectedly, analysis of our model shows that, by alternating between these two losing strategies, *P*_1_ outcompetes *P*_2_. This counter-intuitive result recapitulates a phenomenon that is known as Parrondo’s paradox in game theory (Harmer and Abbott, 1999). Thus, alternating between lysis and lysogeny appears to be intrinsically beneficial for a phage.

## Results

### Alternating between losing lysis and lysogeny strategies results in a winning strategy for a phage

A mathematical model was designed to measure the outcomes of infection between two competing phages that differ in their rates of adsorption, mortality, burst size (i.e., number of progeny virions per infection), and, principally, their propensities for lysis or lysogeny. One of the key parameters of infection driving viral replication strategies is host cell density (Knowles et al., 2016; Thingstad, 2000; Weitz et al., 2015). Thus, to realistically capture phage lysis-lysogeny switches, the decisions are not represented by a constant but rather by a probability function *μ*(*h*), where *h* represents the density of host cells (Equation 3, see Methods and Supplementary Information for details on *μ*(*h*)). A detailed list of all parameters incorporated in the model, their descriptions and initial conditions are given in Supplementary Table S1. In all competitions, the first phage (*P*_1_) is temperate and set at a complete disadvantage to the second phage (*P*_2_), regardless of whether *P*_1_ replicates using the lytic or lysogenic pathway. Specifically, *P*_1_ is penalized with a smaller burst size (*ρ*), lower infection rate (*η*(*h*)) and higher mortality rate during lysogeny (*d_l_*) or lysis (*d_v_*).

In the first competition, the lysis-lysogeny probability function (Equation 3) was set to zero for both phages, which means that all cells are lysogenized upon infection by either phage. As expected, *P*_1_ immediately becomes extinct and *P*_2_ dominates the environment (losing strategy I, **Fig. 1A, 1C**). We note that both the lytic (*v*_2_) and lysogenic (*l_2_*) stages of phage *P*_2_ development are present, reflecting induction (Boling et al., 2020), whereby a fraction of *P*_2_ lysogens transition to the lytic pathway. A similar result is obtained in the next competition where both phages instead enter the lytic pathway upon infecting the host cells, which is achieved by setting the lower bound *α* and range *β* of the probability function (*α,β*) = (1,0). In this competition, *P*_2_ infects all susceptible host cells and drives *P*_1_ to extinction (losing strategy II, **Fig. 1A, 1C**). Thus, neither the purely lytic nor the purely lysogenic pathways are viable strategies for *P*_1_.

**Fig. 1.**
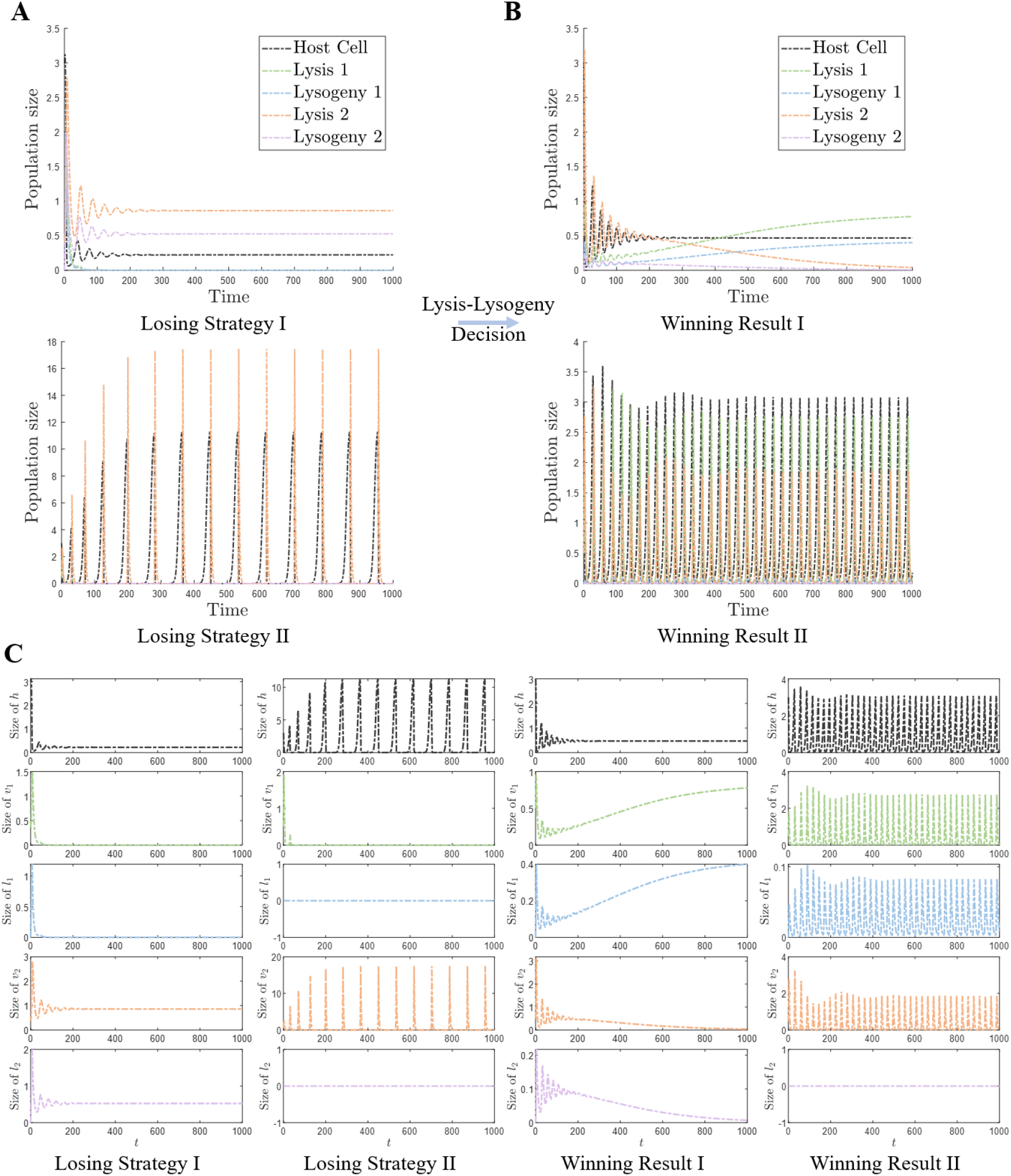
Competition between two phages with different properties. The parameter values are from initial condition 1 in Supplementary Table S1 unless stated otherwise. **a,** Two “losing” strategies of a disadvantaged phage (*P*_1_), in competition with an advantaged phage (*P*_2_). All infected cells become lysogenic in the losing strategy I, and all infected cells are lysed in the losing strategy II. **b,** Two “winning” outcomes reached by *P*_1_. Winning outcome I is reached in competition with a temperate phage *P*_2_, and winning result II is obtained in competition with a virulent phage *P*_2_. **c,** Population dynamics in different competitions.

In the next competition, both phages switch between the lytic and lysogenic pathways. If *P*_1_ has a higher propensity for lysogeny than *P*_2_ (initial condition 1, see Supplementary Information), *P*_1_ becomes dominant at a time point between 300 and 500 (**Fig. 1B-C**, winning result I), with the lytic form of *P*_1_ (*v*_1_) overtaking the lytic form of *P*_2_ (*v*_2_) at around *t* = 300 and gradually driving *P*_2_ to extinction. The ability of *P*_1_ to drive *P*_2_ to extinction is attributable to the rise of *P*_1_ lysogens (*l*_1_), reflecting the importance of the lysogenic pathway for a disadvantaged phage to outcompete an advantaged phage under these conditions.

We next competed *P*_1_, which alternated between the lytic and lysogenic pathways, against a purely lytic phage *P*_2_, to determine if and when a winning outcome by *P*_1_ could be achieved. Specifically, *P*_2_ replicates through a purely lytic pathway by setting (*α, β*) equal to (1,0) in the probability function *μ*(*h*). For a fair comparison, in this experiment (*α, β*) for *P*_1_ is set as (0.9,0.1), so that most but not all infections by *P*_1_ follow the lytic pathway. Under this initial condition, *P*_1_ overtakes *P*_2_ immediately after the first wave of growth around *t* = 100 (**Fig. 1B-C**, winning result II), which is faster than the winning outcome reached by *P*_1_ in competition against another temperate phage (winning result I). Unlike the previous experiment where *P*_2_ was driven to extinction, *P*_1_ and *P*_2_ coexist in the environment, but *P*_1_ reaches a much higher abundance. This result supports the results of the preceding experiment demonstrating that alternating the lytic strategy with the lysogenic one is essential for a disadvantaged phage to outcompete a phage with more favorable life history traits that employs a single strategy, in this case, the lytic one.

The two winning strategies employed by *P*_1_ yielded contrasting results for the competitor, namely, *P*_2_ was either driven to extinction or persisted stably at a lower relative abundance. Therefore, we examined the range of lysis-lysogeny decisions by *P*_1_ that allow *P*_2_ to coexist over a longer timescale. The timescale of the experiment was increased fivefold relative to that in **Fig. 1** and the lysis-lysogeny propensities from winning result II (*α,β*) = (0.9,0.1) were used as a control. As expected, the same outcome from winning result II was achieved when increasing the timescale, demonstrating a long-term coexistence between the winning phage *P*_1_ and the losing competitor *P*_2_ (**Fig. 2A**). If the probability of *P*_1_ to lysogenize is increased (*α*_1_ = 0.7), *P*_1_ wins the competition without driving *P*_2_ to extinction (**Fig. 2B**). Interestingly, within the time intervals *t* ∈ [1200,2000] and *t* ∈ [2800,3200], *P*_2_ temporarily regains dominance over *P*_1_ (**Fig 2B**), because *P*_2_ infects host cells without restriction by adopting solely the lytic pathway. However, *P*_1_ reaches a much higher abundance than *P*_2_ in the course of the long-term coexistence by alternating lytic and lysogenic pathways. If the probability for *P*_1_ to lysogenize is increased further (*α*_1_, = 0.5), *P*_2_ is immediately driven to extinction after the first wave (**Fig. 2C**). Collectively, these results emphasize the role of the lysis-lysogeny switch as the principal life history trait mediating inter-viral conflicts.

**Fig. 2.**
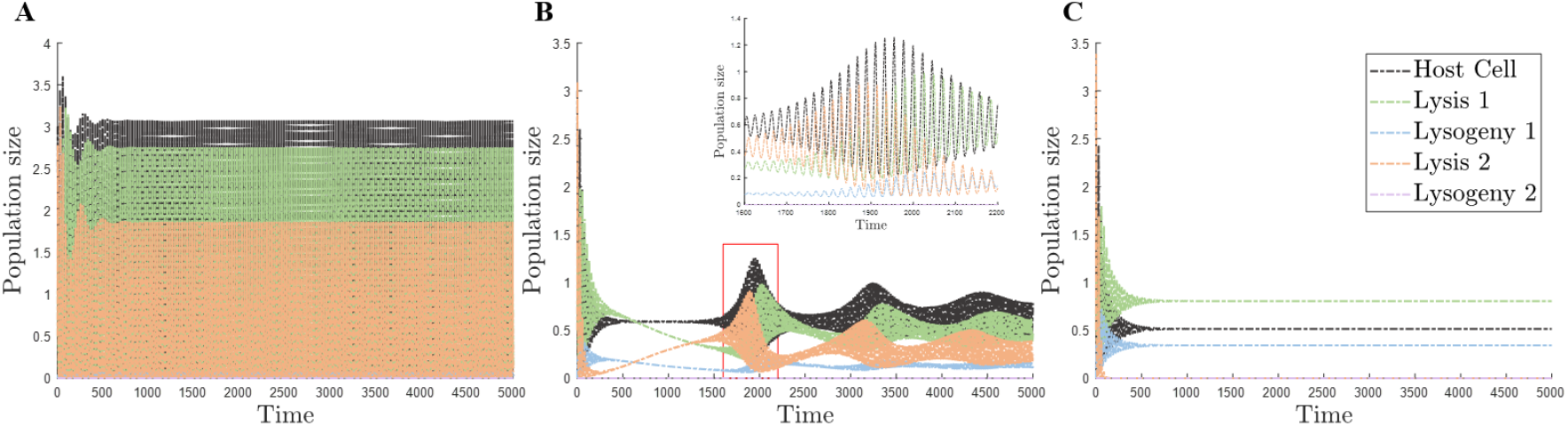
Long-term competition between a virulent phage and a temperate phage with different values of *α*. The lower bound *α* of the probability function gradually decreases, which means a higher probability to lysogenize. The parameter values are from the initial condition 1 as in Supplementary Table S1 unless stated otherwise. **a,** The parameters are the same as the parameters in the winning result II of Fig. 1B, so that the two phages coexist long-term. **b,** The lower bound *α*_1_ equals 0.7 resulting in a long-term coexistence between the two phages. An important transformation is shown in the inset. **c,** The lower bound *α*_1_ equals 0.5, in which case *P*_2_ goes extinct.

### Combinations of model parameters yielding Parrondo’s paradox

We then ran a comprehensive simulation to determine the range of parameters that support winning outcomes for *P*_1_, rather than using a single parameter set. The outcomes were evaluated by averaging the abundance of each competitor’s lytic and lysogenic progeny when the competition reached steady state (between *t* = 2900 and *t* = 3000). Because an all-vs-all parameter comparison would be impractical to examine, we present the results of this systematic survey by considering three qualitatively categorized sets of parameters: environmental (carrying capacity *K*, mortality rates of the virus *d_v_* and of the host *d_h_*), lysis-lysogeny switch (switching interval, switching threshold, lower and upper bounds), and phage life history traits (burst size *ρ*, infection rate *η*(*h*), and induction rate *γ*). By exploring the range of parameters and evaluating the outcome of the competition, we can determine whether Parrondo’s paradox is observed over a relatively narrow or over a broad range of conditions.

We first explored the space of environmental parameters, the carrying capacity *K*, host cell mortality rate *d_h_* and phage mortality rate *d_v_* parameter spaces. The carrying capacity *K* limits the total number of cells in the environment, including both uninfected hosts and lysogens emerging upon infection by either phage. *P*_1_ reaches higher abundance compared to *P*_2_ when the carrying capacity is relatively large, but the opposite is true when the carrying capacity is small (**Fig. 3A**), indicating that the total number of cells in the environment affects the ability of a disadvantaged temperate phage to outcompete an advantaged phage. Both phages increase in abundance when the host cell mortality rate decreases (**Fig. 3A**), reflecting the shared ‘interests’ of integrated phages and their host cells (Argov et al., 2017). Evidently, decreasing the mortality rate of *P*_1_ (*d*_*v*_1__) (**Fig. 3B**) or increasing the morality rate of *P*_2_ (*d*_*v*_2__) (**Fig. 3C**) buttresses the winning outcome for *P*_1_. When *d*_*v*_1__ and *d*_*v*_2__ are both changed at the same time, there is still a well-defined dividing region in which *P*_1_ transitions from a loser to a winner (**Fig. 3D**). In this range, *K* exerts no obvious effect on the competition result (**Fig. 3B-C**) because the total number of cells is relatively small and therefore the growth rates of uninfected hosts (*h*) and lysogens (*l*) do not change drastically. Thus, the mortality rates have a greater impact on the outcome of the competition compared to the environmental carrying capacity.

**Fig. 3.**
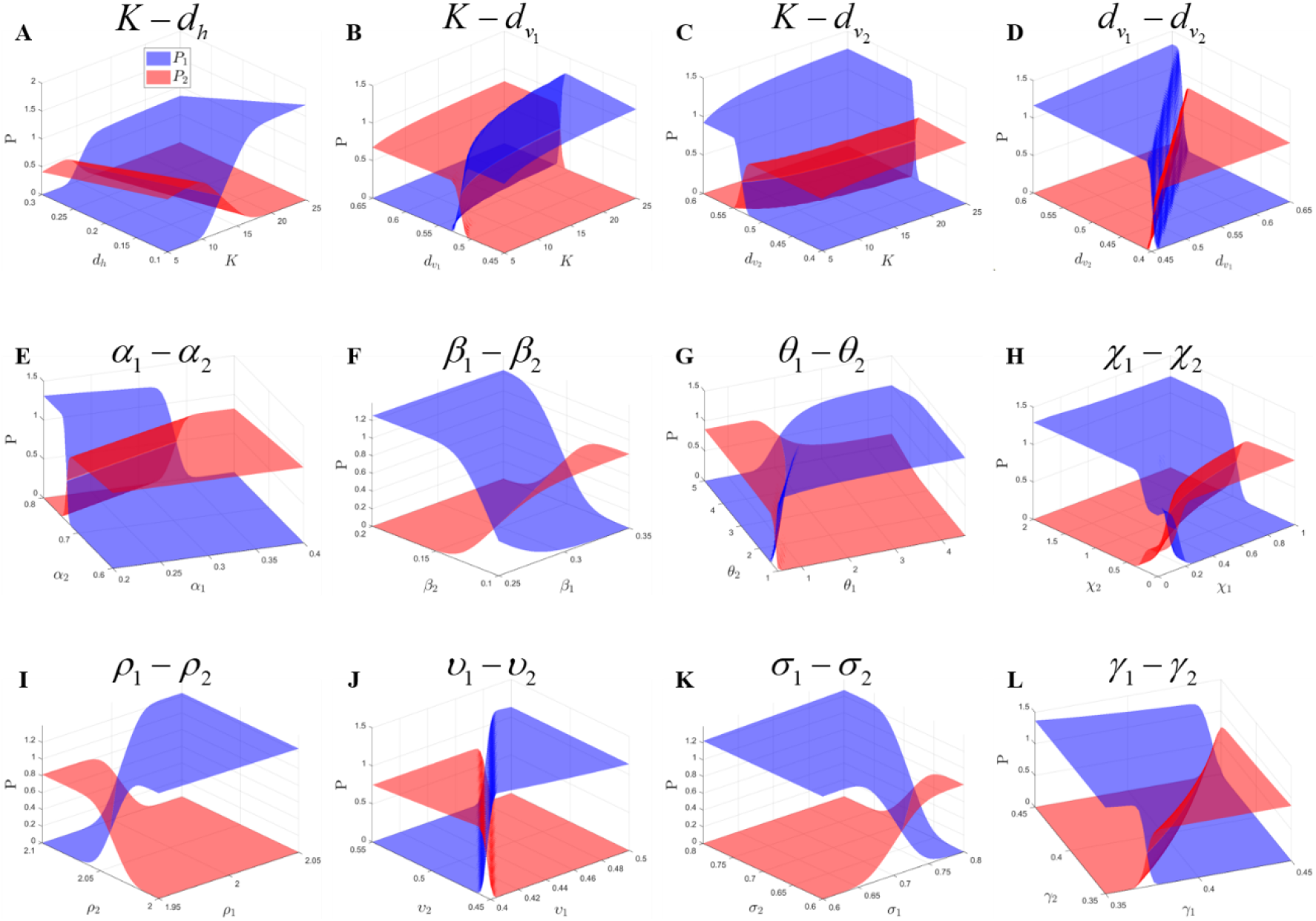
The regions of the parameter spaces yielding Parrondo’s paradox in the competition between two phages. The parameter values are from initial condition 1 in Supplementary Table S1 unless stated otherwise. **a-d,** The change in environment carrying capacity *K* is compared with the mortality rate of host cell and virus in lytic phase *d_h_* and *d_v_*, demonstrating the mortality rates have more influence on the winning result compared with the change of *K*. **e-h,** The parameters of the probability function between two phages are compared, including *α, β, θ*, and *χ*. **i-l,** The key properties of two phages are compared, including *υ* and *σ* in infection rate function, as well as the burst size *ρ* and the induction rate *γ*.

The probability function *μ*(*h*) that models the phage lysis-lysogeny switch was demonstrated to substantially impact the outcome of the competition (**Fig. 2A-C**). Thus, the effect of each of the four parameters of this function was evaluated across a wide range of values (**Fig. 3E-H**). The two parameters with the greatest influence on the outcome of the competition in our initial experiments (**Figs. 1-2**) were *α* and *β*, which set the lower and upper bounds of the probability of a phage to enter the lytic pathway, respectively. A phage is driven towards the lytic pathway by increase of *α* and decrease of *β* (**Fig. 4A**). Comparison of the *α* values between the two phages shows that *P*_1_ outcompetes *P*_2_ in a narrow range where the *α* value of *P*_1_ is approximately half that of *P*_2_ (**Fig. 3E**), or in a broad range where *β*, the value of *P*_2_ is greater than ~0.15, (**Fig. 3F**). *P*_1_ can also win the competition by changing the *α* and *β* values while competing against a temperate (**Fig. 4B**) or purely lytic (**Fig. 4C**) competitor with fixed *α* and *β* values. With regard to the other parameters related to the lysis-lysogeny switch, *P*_1_ can dominate over relatively broad parameter spaces. *θ* is the switch threshold, so that a large *θ* causes a phage to switch from lysis to lysogeny in the presence of a relatively large number of hosts (**Fig. 3G**). The parameter *χ* is the switching interval, with a small value of *χ* enabling quick transition to lysogeny with the decline of *h* (**Fig. 3H**). Collectively, these results indicate that a disadvantaged phage can win the competition by optimizing multiple parameters involved in switching between lysis and lysogeny.

**Fig. 4.**
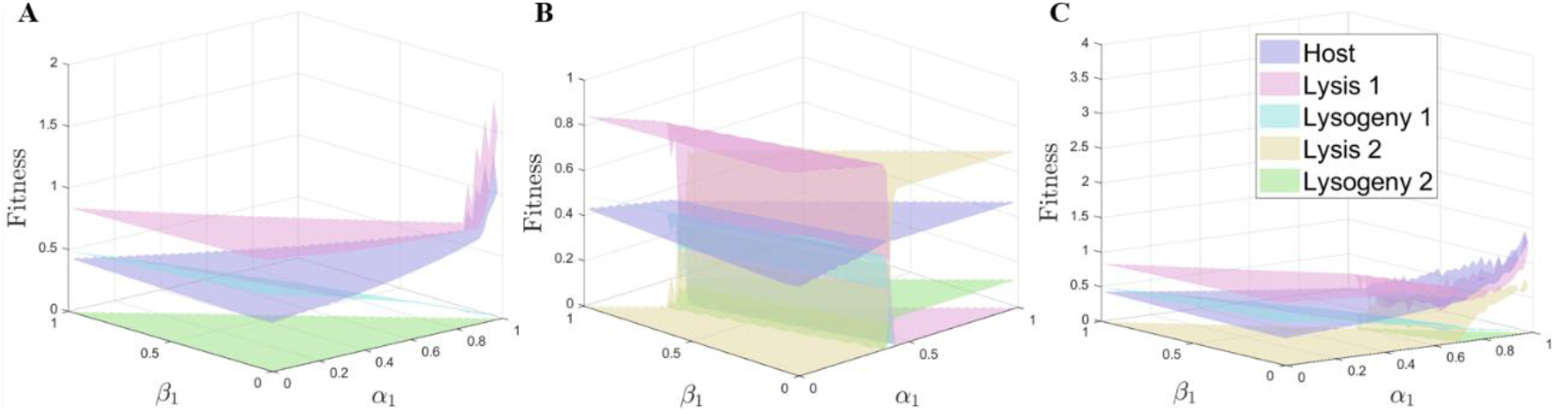
Fitness of two phages and the host depending on the parameter values. Fitness is defined as the mean size of a population at steady state (*t* ∈ [2900,3000]). The parameter values are from the initial condition 1 in Supplementary Table S1 unless stated otherwise. **a,** Fitness of each population in the presence of *P*_1_ alone. **b,** Fitness of each population under the competition between two temperate phages ((*α*_2_, *β*_2_) = (0.8,0.2)). **c,** Fitness of each population under the competition between a temperate phage and a virulent phage ((*α*_2_, *β*_2_) = (1,0)).

We next examined the life history traits that are traditionally analyzed in models of viral fitness, including burst size (*ρ*), infection rate (*η*(*h*)) and induction rate (*γ*).

Evidently, a large burst size results in a greater number of infectious progeny viruses, so that the phage with the larger burst size will win the competition against th one with the smaller burst size (**Fig. 3I**). The infection rate *η*(*h*) of each phage is determined by the efficiency *υ* and handling time *σ* (Equation 2, see Methods and Supplementary Information for details on *η*(*h*)). Increasing the overall infection rate with larger values of *υ* or smaller values of *σ* enable a phage to infect more susceptible cells than the competitor (**Fig. 3J-K**). Not surprisingly, there are more lysogens when the induction rate *γ* is low, and the existence of lysogens affects the outcome of the competition (**Fig. 3L**). Although either phage can reach a higher abundance than the other by optimizing these traits, these results show that the steady state number of *P*_1_ winners is larger than the steady state number of *P*_2_ winners.

The impact of the range of lysis-lysogeny decisions on the outcome of the competition was further studied in terms of the fitness of the host and phage populations under different probabilities of cell to lysis (**Fig. 4**). Fitness is defined as the mean size of a population at steady state (*t* ∈ [2900,3000]),showing the dynamics of all populations, both host and the two phages (*P*_1_ or *P*_2_) under different lysis-lysogeny decisions and thus providing additional insight into the lysis-lysogeny decision. Specifically, the lower bound (*α*_1_ and the upper bound *α*_1_ + *β*_1_ follow the inequalities 0 ≤ *α*_1_ ≤*α*_1_ + *β*_1_ ≤ 1. In **Fig. 4A**, the host is only infected by *P*_1_, and the fitness of *v*_1_ is higher than the fitness of *h* in the corresponding region of the parameter space. The fitness of *l*_1_ is the lowest among these three populations and declines with the rise of *α*_1_. Moreover, the lytic pathway is favored by the phage with the increase of *α*_1_ and the decrease of *β*_1_ (**Fig. 4A**). In the competition between two temperate phages, *v*_1_ has greater fitness when *α*_1_ is small (and *β*_1_ is large), and *v*_2_ has greater fitness when *α*_1_ is large (and *β*_1_ is small), likely, because the lower value of *α*_1_ renders more phage in the lysogenic phase under environmental fluctuation, resulting in high fitness of *P*_1_ (**Fig. 4B**). This effect is crucial for the phage in an adverse environment. The fitness of both phages is greater than the host fitness, and the phage in the lytic phase is fitter than the phage in the lysogenic phase (**Fig. 4B**). In the competition between the virulent phage and temperate phage (**Fig. 4C**), the fitness of *v*_1_ is also greater than the fitness of *h* akin to the result in **Fig. 4A**.

## Discussion

We demonstrate here that a phage with inferior life history traits can outcompete a competitor with superior traits over a broad set of conditions by switching between the lysis and lysogenic infection strategies. Neither the lytic nor the lysogenic phase of the disadvantaged phage is competitive on its own, yet a winning outcome is achieved by alternating between the two strategies (**Fig. 1**). This counterintuitive result is formulated as Parrondo’s paradox, whereby alternating between two losing strategies can result in a winning outcome. This effect is manifested at different levels of biological organization (Cheong et al., 2019), such as the alternation between unicellular and multicellular phases in organismal life history (Cheong et al., 2018), between phases of activity and dormancy in predator and prey (Tan et al., 2020) or between nomadic and colonial life styles (Tan and Cheong, 2017).

Several studies have examined the conditions in which lysogeny is more favorable than lysis for phages. Vacillations in host cell population density, due to environmental downturns or other factors, could lead to extended periods of low host availability that are too low to support lytic infection (Stewart and Levin, 1984). Lysogenizing host cells under these conditions allays the risk of population collapse (Maslov and Sneppen, 2015). As the host cell population recovers, this strategy of leveraging binary cellular fission for vertical transmission becomes inferior to horizontal transmission (that is, lytic reproduction) (Li et al., 2020; Weitz et al., 2019). Consistent with lysogeny being favorable at low host cell availability, our experiments indicate that, when the number of host cells oscillates, temperate phages that favor the lysogenic phase persist, whereas those that prefer lysis as well as purely lytic phages go extinct (**Fig. 1-2**).

Fewer studies have investigated the parameters of infection that dictate the outcome between two phages competing for a host cell. The ubiquity and biological importance of inter-virus competition is perhaps best reflected by the broadly distributed and diverse superinfection exclusion mechanisms that prevent secondary infections of lysogens by other phages (Dedrick et al., 2017; Gentile et al., 2019; Mavrich and Hatfull, 2019). Mathematical models predict that temperate phages can invade microbial populations in the presence of a competing lytic virus, so long as they confer a minimal level of superinfection immunity (Li et al., 2020). In one of our experiments, the disadvantaged phage indeed confers superinfection immunity to the host cell during lysogeny and drives the competitor to extinction (**Fig. 2**). With the decrease in the probability to lysogenize, the disadvantaged phage is instead driven to extinction. Notably, we identified multiple regions of the parameter space where the two phages can coexist, with the disadvantaged phage that alternates between lysis and lysogeny reaching higher abundance (**Figs. 3,4**).

Our work identifies the conditions in which a disadvantaged phage can not only coexist with but outcompete a phage with superior life history traits by cycling between two losing strategies, a clear case of Parrondo’s paradox. Parrondo’s paradox applies to large regions of the parameter space, with propensities for lysis or lysogeny being the primary parameters of infection that determine this winning outcome (**Fig. 2**). It should be emphasized, however, that many combinations of parameters favor a pure lytic or lysogenic strategy.

On closer examination, the advantage of alternating modes of interaction with the host by a virus does not appear truly paradoxical, being tightly linked with the oscillations of the host population that are well known from the classic, Lotka-Volterra-type prey-predator models (Roughgarden, 1979). Indeed, once the host cell density drops, due to the cell lysis by the virus, the latter switches to lysogeny, but when the host population recovers, switching to lysis becomes advantageous. Combined with superinfection exclusion, such biphasic lifestyle seems to be optimal for viruses within a broad range of infection and environmental parameters. Analysis of the model described here shows that the advantage of such flexibility is high enough to overcome imperfection in each of the individual strategies.

Lysogeny likely evolved as an adaptation to host population oscillations caused by the virus itself or by other environmental factors. Given that such oscillations are a generic feature of population dynamics, Parrondo’s paradox is likely to be a common if not universal phenomenon in host-parasite interactions.

## Acknowledgements

K. H. C. and T.W. are supported by the Singapore University of Technology and Design (Grant No. SRG SCI 2019 142). S.B. and E.V.K. are supported by the Intramural Research Program of the National Institutes of Health of the USA (National Library of Medicine).

## Author Contributions

K. H. C. and E.V.K. designed research; T. W. and K. H. C. performed research; T. W., K. H. C., S.B. and E.V.K. analyzed the data. K. H. C., T. W., S.B. and E.V.K. wrote the manuscript.

## Methods

### Dynamic Model of Lysis-Lysogeny

#### Evolution Model

The dynamic evolution model includes two phages (*P*_1_ and *P*_2_) competing for a sensitive host cell (*h*). Upon infection of the host, the progeny of either phage can be lytic (*V*) or lysogenic (*l*) (the subscripts of all variables described refer to phages *P*_1_ or *P*_2_). Host cells lysogenized by a phage are immune to infection by the competitor phage (i.e., *P*_1_ prevents superinfection of the host cell from *P*_2_ and vice versa). The environment carrying capacity (*K*) determines the growth rate of the host cell and the lysogenic virus, with the maximum growth rate set by *τ*, which follows the logistic growth model. A susceptible host cell can be infected by two phages with different rates *η*(*h*), which are determined by the density of the host cells. Host cell death is quantified by the rate *d_h_* and is supplemented by the loss of prophage-mediated immunity to the competing virus with the rate *λ*. Upon infection of a host cell, lysis probability is quantified by *μ*(*h*), and lysogeny probability is 1 — *μ*(*h*), where the probability is determined by *h*. A phage in the lysogenic phase (prophage) can be induced to enter lytic phase at the rate *γ*. Both infection and induction in the lytic phase have the burst size (reproduction rate) *ρ*. Free virions are inactivated at the rate *d_v_* and prophages *l* are inactivated at the rate *d_l_*. Thus, the model is captured by the following differential equations,

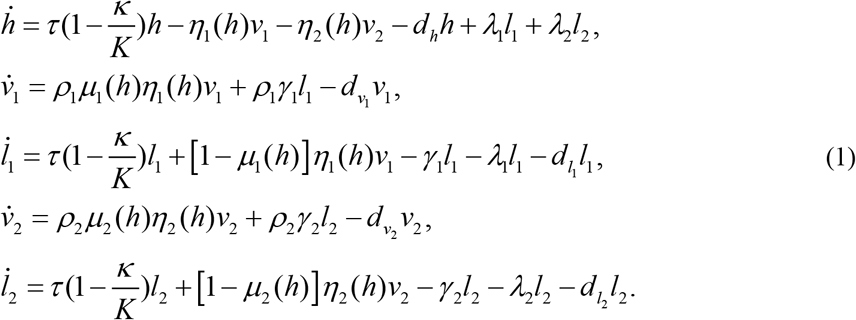

where *K* is the total number of cells, which equals *h* + *l*_1_ +*l*_2_. The flow and interaction among these compartments are shown in Supplementary Figure S1, and the initial values for each compartment and parameters are shown in the initial condition 1 of Supplementary Table S1. Values for each phage parameter were collected from recent analyses of phage lysis-lysogeny decisions (Pleška et al., 2018; Sinha et al., 2017; Wahl et al., 2019).

#### Parameter Functions

The infection function *η*(*h*) is modelled by the Holling type II functional response (Rosenzweig and MacArthur, 1963), which is monotonically increasing. This function takes the form

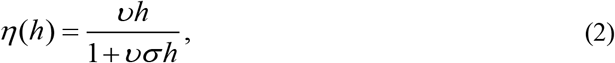

where *υ* is the efficiency, and *σ* is the handling time that phages need to infect the host cell. This function is linear: *η*(*h*) ≈ *υh* when *h* → 0, and equals to 1/ *σ* when *h* →∞.

The probability function *μ*(*h*) for host cell lysis is determined by the modified sigmoid function (Kuwamura et al., 2009),

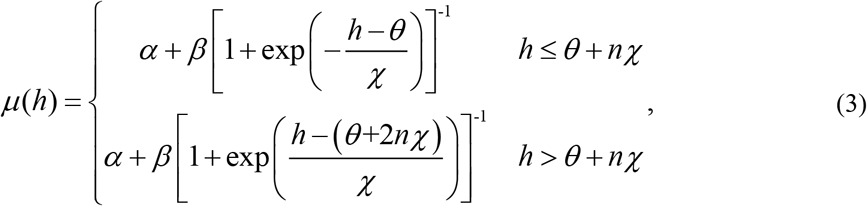

where *α* is the lower bound, and *β* is the range which determines the upper bound (*α + β*). *θ* is the switching threshold *μ*(*θ*) = *α* + *β* / 2. *χ* determines the shape of the function, whereby sharper switching occurs at smaller *χ. θ* + *nχ* is the turning point of the trend, and *n* = 2 (Supplementary Figure S2) shows two diagrams of *η*(*h*) and *μ*(*h*) with their key properties.

## Supplementary Information

Our source code can be accessed at: https://osf.io/vth96/?view_only=c61c0a312ef04624acbeb41d071e70df

Table S1. The parameters of the model and their initial values.

Figure S1. The framework of the model showing the relationship between the compartments.

Figure S2. Diagrams of *μ*(*h*) and *η*(*h*).

Figure S3. Competition between two phages with different properties (based on initial condition 2 in Table S1).

## Supplementary Information

**Table S1.** The initial values and descriptions of parameters

| Parameter | Description | Initial condition 1 |  | Initial condition 2 |  |
| --- | --- | --- | --- | --- | --- |
| $h$ | Number of host cells | 3 | | 4 | |
| $v_1$ | Number of the first virus particles in lytic phase | 0.5 | | 2 | |
| $v_2$ | Number of the first virus particles in lysogenic phase | 0 | | 0 | |
| $l_1$ | Number of the second virus particles in lytic phase | 0.5 | | 2 | |
| $l_2$ | Number of the second virus particles in lysogenic phase | 0 | | 0 | |
| $\tau$ | Growth rate | 0.5 | | 0.5 | |
| $\kappa$ | Total number of cells | $h + l_1 + l_2$ | | $h + l_1 + l_2$ | |
| $K$ | Environment carrying capacity | 20 | | 30 | |
| $d_h$ | Mortality rate of the host cells | 0.2 | | 0.25 | |
| | | $P_1$ | $P_2$ | $P_1$ | $P_2$ |
| $\lambda$ | Immune loss rate | 0.05 | 0.05 | 0.05 | 0.05 |
| $\rho$ | Burst size (Reproduction rate) | 2 | 2.05 | 3 | 3.05 |
| $\gamma$ | Induction rate | 0.4 | 0.4 | 0.4 | 0.4 |
| $d_v$ | Mortality rate of virus in lytic phase | 0.55 | 0.5 | 0.45 | 0.4 |
| $d_l$ | Mortality rate of virus in lysogenic phase | 0.25 | 0.2 | 0.2 | 0.15 |
| $\eta(h)$ | Infection rate | Function | Function | Function | Function |
| $\mu(h)$ | Probability to lyse | Function | Function | Function | Function |
| $\upsilon$ | Viral efficiency to infect | 0.45 | 0.5 | 0.45 | 0.5 |
| $\sigma$ | Handling time | 0.7 | 0.7 | 0.6 | 0.6 |
| $\theta$ | Switching threshold | 1 | 1.5 | 1 | 1.5 |
| $\chi$ | Switching interval | 0.5 | 1 | 0.5 | 1 |
| $\alpha$ | Lower bound | 0.3 | 0.8 | 0.3 | 0.8 |
| $\beta$ | Range | 0.3 | 0.2 | 0.3 | 0.2 |

**Figure S1.**
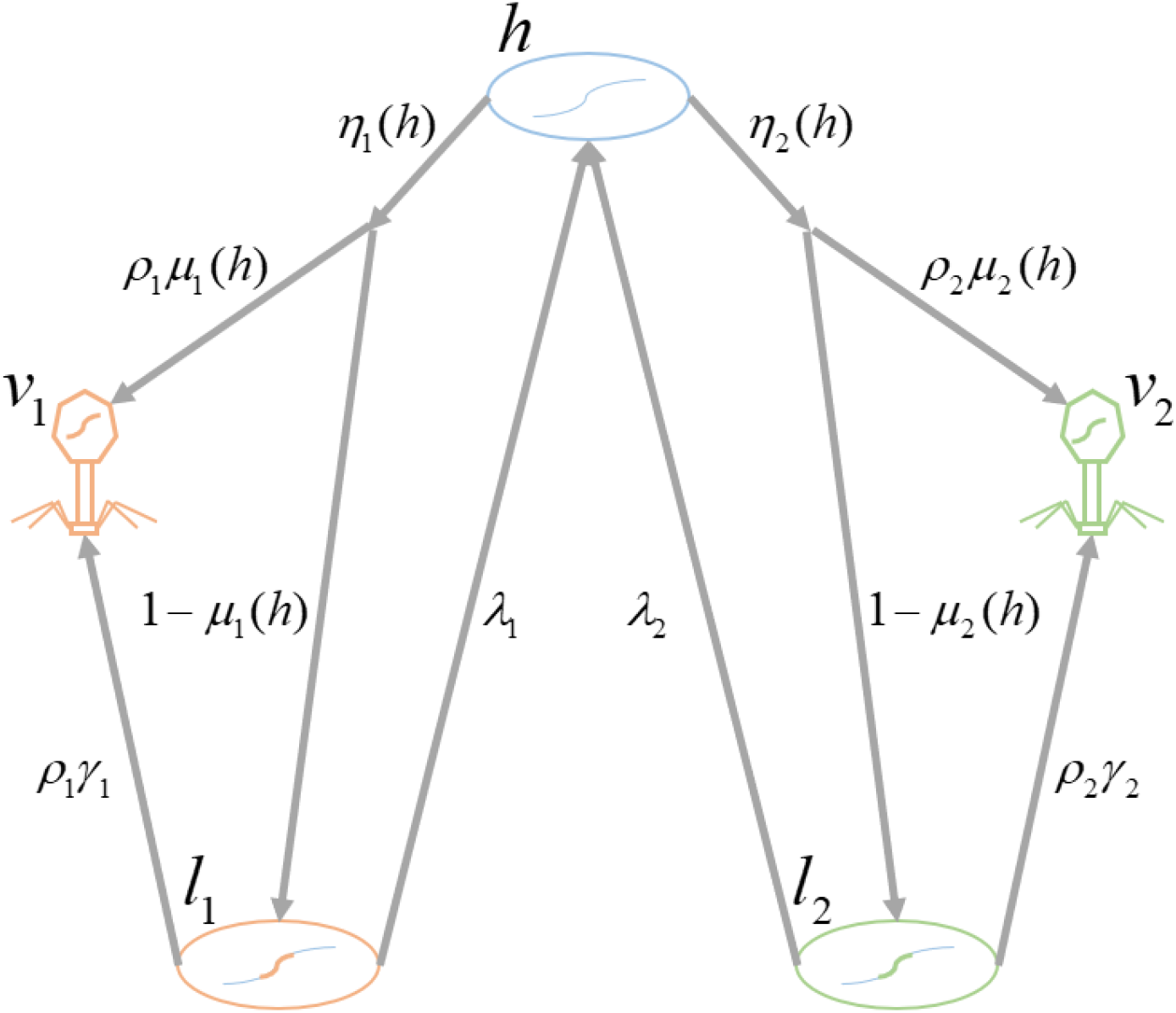
The framework of the model showing the relationship between the compartments.

**Figure S2.**
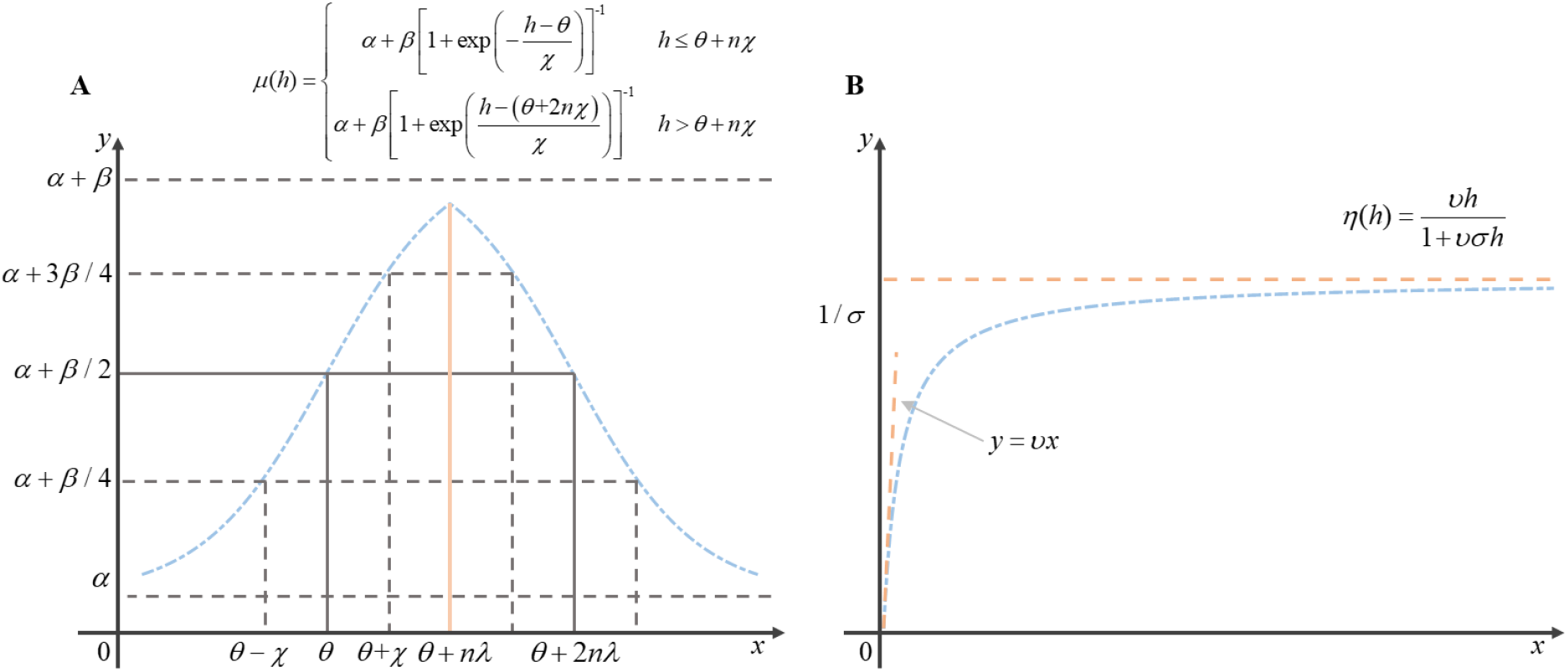
The diagrams of *μ*(*h*) and *η*(*h*). **a**, The diagram of infection function *μ*(*h*) with key points shown in the curve. **b**, The diagram of probability function *η*(*h*), and *η*(*h*) ≈ *υh* when *h* → 0, as well as *η*(*h*) → 1/ *σ* when *h* →∞.

**Figure S3.**
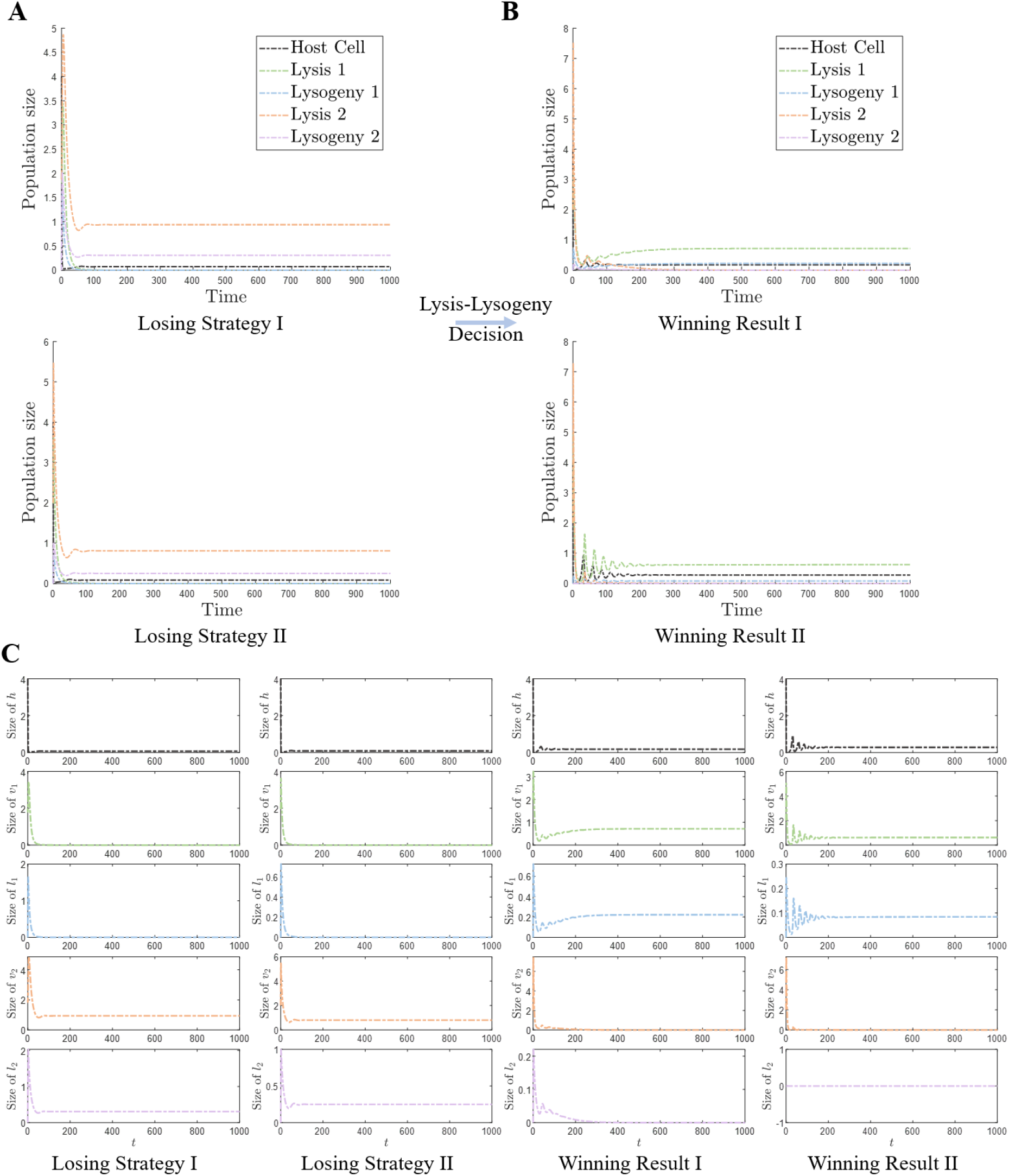
Competition between two phages with different properties. The results are based on the initial condition 2 in Table S1 **a,** Two losing strategies in the competition. All infected cells become lysogeny in the losing strategy I, and all infected cells become lysis in the losing strategy II. **b,** Two winning results between different phages’ competition. winning result I is obtained by the competition between two temperate phages, and winning result II is obtained by the competition between temperate phage and virulent phage (*α*_1_,*β*_1_) = (0.8,0.1)). **c,** The specific population changes based on different competitions.

